# The Underground World of Plant Disease: How Does Rhizosphere Dysbiosis Affect Plant Health Above-ground?

**DOI:** 10.1101/2024.02.27.582369

**Authors:** Toi Ketehouli, Josephine Pasche, Victor Hugo Buttrós, Erica M. Goss, Samuel J. Martins

## Abstract

Similar to the human gut microbiome, diverse microbes colonize the plant rhizosphere, and an imbalance of this microbial community, known as dysbiosis, may negatively impact plant health. This study aimed to investigate the influence of rhizosphere dysbiosis on above-ground plant health using tomato plants (*Solanum lycopersicum* L.) and the foliar bacterial spot pathogen *Xanthomonas perforans* as model organisms. Four-week-old tomato plant’s rhizospheres were treated with streptomycin (0.6 g × L^-1^), or water (negative control) and spray-inoculated with *X. perforans* (10^5^ cells × mL^-1^) after 24 h. Half of the plants treated with streptomycin and *X. perforans* received soil microbiome transplants (SMT) from uninfected plant donors 48 h after streptomycin application. Streptomycin-treated plants showed a 26% increase in disease severity compared to plants that received no antibiotic, while plants that received the SMT had an intermediate level of disease severity. Antibiotic-treated plants showed a reduced abundance of rhizobacterial taxa like Cyanobacteria from the genus *Cylindrospermum* as well as down-regulation of genes related to plant primary and secondary metabolism and up-regulation of plant defense genes associated with induced systemic resistance (ISR). This study highlights the crucial role of beneficial rhizosphere microbes in disease resistance, even to foliar pathogens.

## INTRODUCTION

Plants and microbes have developed a close symbiotic relationship estimated to date back hundreds of millions of years (Martin et al 2016). As a result of this relationship, plants and microbes have evolved very sophisticated beneficial communication. For instance, under stress (e.g., a pathogen infection), plants change their root chemical composition by releasing modified root exudates and “*cry for help*” to recruit beneficial microbes from the soil to cope with the stress (Martins et al 2023, Rolfe et al 2019). Like the human gut microbiome, the plant rhizosphere, the narrow soil area around the root, is colonized by diverse microbes (Ravelo-Ortega et al 2023). Some microbes play a crucial role in plant physiological processes and are a first line of defense against invading pathogens (Berg et al 2020, Jiang et al 2022, Trivedi et al 2020). Plant defense mediated by the rhizosphere microbial community can happen directly, through microbe-pathogen inhibition or indirectly, via induced systemic resistance (ISR) (Mehmood et al 2023, Oleńska et al 2020, Salwan et al 2023). An imbalance of the microbial community, known as dysbiosis, can influence plant fitness. For instance, rhizospheric dysbiosis in tomato plants, evidenced by a disruption in Firmicutes and Actinobacteria population, led to a 1.5-fold increase for the below-ground bacterial wilt caused by *Ralstonia solanacearum* and also a significant increase in *Fusarium oxysporum* wilt (Lee et al 2021, Zhang et al 2021). In these examples, the reverse of dysbiosis, also known as eubiosis, leads to the hypothesis that bacteria in the rhizosphere (below-ground) have essential health functions to below-ground plant infections.

In some cases, the soil microbiome transplant (SMT, hereafter) from suppressive soils can cause conductive soils to become suppressive (Schlatter et al 2017, Weller et al 2002) Research into microbiome transplantation has been pioneered and explored in the context of human health, specifically the transfer of fecal gut microbiomes from healthy donors to cancer patients. This concept of SMT has recently been investigated in plant health, where the focus is on transferring a healthy plant microbiome to plants experiencing dysbiosis (Bziuk et al 2022). The process of microbiome transplantation involves the extraction of a healthy donor microbiome and application into a recipient soil, and the addition of the microbiome can then cause disease-suppressive effects (Gowda et al 2023). Understanding the dynamics of plant health/disease processes concerning rhizosphere dysbiosis is essential for efficiently and sustainably managing plant diseases.

Bacterial spot, or bacterial leaf spot, is an economically significant disease of tomato plants (*Solanum lycopersicum* L.), caused by *Xanthomonas* spp. (Strayer et al 2016). Management of bacterial spot disease on tomato is challenging due to limited host resistance, pathogen tolerance to copper bactericides, and the aggressiveness of the pathogen (Abrahamian et al 2019). streptomycin is an antibiotic that is labeled for use on greenhouse tomato production, which can be effective when target strains are sensitive (Abrahamian et al 2019) Since the 1950s, when streptomycin was licensed to combat the bacterial disease fire blight in apple trees, antibiotic use has expanded to other pathosystems, including bacterial spot disease in tomato and pepper plants and “Huanglongbing” on citrus trees (Kirchhelle 2018, Yin et al 2023). Despite its expansion in use, several concerns have been raised with antibiotic application in agriculture, such as the increase of antibiotic-resistant bacterial pathogens (Stockwell and Duffy 2012) in the environment and the impact on non-target pollinators and microbes (Chen et al 2019), contributing to the vacate EPA exemption of Streptomycin use in Florida’s Citrus orchards since December of 2023 (EPA 2023). Streptomycin, along with other antibiotics, are still used in the Americas and Asia, but banned in Europe. Its usage continues in certain areas, even against local regulations, especially in low- and middle-income countries. Gathering detailed information about each country’s specific laws on antibiotic use in agriculture proves difficult, as the legislation is typically available only in the local languages (Miller et al 2022, Verhaegen et al 2023).

In this study, we hypothesized that the induction of dysbiosis in the plant rhizosphere (below-ground, induced by antibiotic application) would influence the plant’s health against the foliar disease bacterial spot (above-ground). The objectives of this study were: [1] investigate the influence of rhizosphere dysbiosis on above-ground plant health using tomato plants and *Xanthomonas perforans* as model organisms; [2] assess the transcriptional and physiological plant responses under dysbiosis; [3] evaluate the effect of soil microbiome transplants (SMT) on the above-ground plant health; [4] assess through microbiome network analysis the possible players in promoting plant health.

## MATERIAL AND METHODS

### Experimental set-up

The soil underwent a comprehensive analysis of the physiochemical parameters, as detailed in Supplementary Table 1. Soil was collected from a field with a history of solanaceous crops (26 m elevation, 29° 38’ 14.064” N, 82° 21’ 21’ 40.6548” W). The soil was blended in a 1:1 ratio with potting soil to enhance moisture retention. Tomato seeds (*Solanum lycopersicum* L. cv Alisia Craig) were grown in 1 L pots, and at 3-weeks-old, 5 seedlings of each treatment were treated with: [1] streptomycin (S); [2] streptomycin followed by addition of microbiome transplant from healthy tomato plants (SMT); [3] water-only (C – negative control). Streptomycin was applied to the soil of each plant using 250 mL of solution at 0.6 g × L^-1^ of antibiotics. Plants were watered daily after inoculation, until field capacity, and kept in the greenhouse of the plant pathology department at the University of Florida (26 m elevation, 29°38’20.238”29°38’20.238” N, 82°21’21.9708”82°21’21.9708” W) under the following conditions: 28°C and relative humidity at 60-65% during all measurements.

### Soil microbiome transplant (SMT)

The extraction of the soil microbiome for transplantation followed the techniques from Silva et al (2022), which were used with some modifications. The soil microbiome from three healthy tomato plants was used to transplant the antibiotic-treated plants. For each recipient plant, 15 g of soil was collected from the root zone of the donor plants and placed in a beaker. Sodium phosphate solution [10 mM] was added to the soil in a 1:10 ratio, and 75 g of soil was collected and mixed with 750 mL of sodium phosphate solution for five repetitions. The mixture was blended for 60 s three times for homogenization. The soil mixture was then transferred to a 1000 mL beaker, placed in an ultrasonic bath, and sonicated for 10 min at room temperature according to Steinauer et al (2023) protocol. The soil suspension was then passed through a 200-mesh (0.07 mm) screen sieve to remove root debris and large soil particles. The resulting soil slurry was diluted with distilled water to a volume of 1250 milliliters. Finally, 250 mL of the slurry solution was applied to each plant 24 h after antibiotic application, gently swirling between applications.

### Assessment of the physiological & plant growth parameters

Physiological parameters were analyzed every seven days after microbiome transplantation through gas exchange and photosynthesis rate measurements. The measurements were made using an infrared gas analyzer (IRGA) LI-COR LI-6800. In parameters: flow, water, and CO_2_ injector: on; humidifier: 20%; CO_2_: 20%; CO_2_-reference: 420 mmol; H_2_O-reference: 65%; desiccant: 100%; relative humidity: 50%; light: 1000 mmol, 9:1 red/blue; temperature −28°C. As a standard, measurements were taken on the third branch emitted by the tomato plants on the terminal leaflet. In cases of damage that prevented the device’s reading or coupling, more distal primary leaflets from the same branch were adopted. At the end of the experiment, plants were collected and root separated from shoot, weighted to obtain the fresh biomass, and dried in oven to obtain the dry biomass.

### Pathogen inoculation and disease evaluation

To obtain pathogen cells for inoculation, a pure colony of *Xanthomonas perforans* was used to initiate a liquid culture in 50 mL of Lysogeny Broth medium (LB) (Bertani 1951) in a shaker incubator at 28°C and at 150 rpm for 24 h. To prepare the cell suspension, the fermented broth was centrifuged for 10 min at 10,000 rpm at room temperature (25°C) and the pellet was suspended in sterile distilled water until an optical density (OD_600_) of 0.1 was reached. The tomato plants were then sprayed with the bacterial suspension until dripping point. The plants were covered with transparent polypropylene bags to ensure humidity for inoculum survival. Five days after pathogen inoculation (DAI), plants were scored every other day for the total foliar bacterial leaf spot disease symptoms caused by streptomycin-resistant *Xanthomonas perforans* strain XP1-6 using the Horsfall-Barratt scale: 0% leaf area affected = 1, 1-2% = 2, 3-6% = 3, 7-12% = 4, 13-24% = 5, 25-50% = 6, 51-75% = 7, 76-87% = 8, 88-93% = 9, 94-97% = 10, 98-99% = 11, and 100% dead tissue = 12 (Horskfall 1945). The data was used to calculate the area under the disease progress curve (AUDPC), according to Shaner and Finney (1977).

### DNA extraction

At 21 days after the pathogen inoculation, 5 g of the distal root portion of each plant was collected and placed into a 50 mL falcon tube with 25 mL potassium phosphate buffer. For the rhizosphere extraction, the falcon tubes were vortexed for 30 seconds, then placed in an ultrasonic bath and sonicated for 10 mins at room temperature according to the protocol by Steinauer et al (2023). After sonication, the roots were removed from the tubes, the falcon tubes were centrifuged at (speed), and the supernatant was discarded. A 200 mg soil sample from each falcon tube was taken for DNA extraction using the Zymo Quick DNA Fecal/Soil Microbe MiniPrep Kit (Zymo Research). Before sequencing, DNA purity and concentration were assessed using spectrophotometry (NanoDrop; Thermo Fisher). The DNA was sent for sequencing at SeqCenter in Pittsburgh, PA. The rRNA 16S gene in the V3-V4 regions was sequenced. After clean-up and normalization, the samples were sequenced on a V3 MiSeq 622 cyc flow cell generating 2 × 301 bp PE reads. Raw read data were submitted to the NCBI SRA under the accession number PRJNA1046511.

### Sampling for transcriptome analysis

A transcriptome analysis was performed to elucidate the impact of antibiotic treatment on the rhizosphere microbiome and the response to *Xanthomonas perforans* infection in tomato plants. Tomato plants were divided into two treatments; one was administered 250 mL of streptomycin solution (0.6 g × L^-1^), while the control received an equal volume of distilled water. After 24 h following the antibiotic application, leaves of all plants were inoculated using a bacterial solution of *X. perforans* suspended in phosphate-buffer saline (PBS) at 1 × 10^5^ cells × mL^-1^. At 19 hours after pathogen inoculation, 3 non-senescent leaves from each of the 3 plants of both treated and control groups were carefully collected and immediately frozen in liquid nitrogen. RNA extraction was performed using the Direct-zol™ RNA Kit from Zymo Research. The mRNA enrichment, fragmentation, adapter ligation, size selection, PCR amplification, and RNA sequencing were carried out at SeqCenter, Pittsburgh, PA (Yang et al 2015). The transcriptome data were then analyzed to identify significantly up-regulated or down-regulated genes in response to the antibiotic treatment compared to the control. Significance was determined using a combination of a *p*-value for a false discovery rate (FDR) test of ≤ 0.05 and a log_2_ fold change (FC) value, with up-regulation defined as log_2_(FC) > 0 and down-regulation as log_2_(FC) < 0 (Storey and Tibshirani 2003).

### Gene annotation and functional clustering

Differentially expressed genes were grouped using the functional grouping function on the DAVID platform (Sherman et al 2022). The other genes were manually grouped based on literature and data available in the UniProt (www.uniprot.org). The genes were grouped into five categories: (1) Unknown function for genes with functions yet to be described; (2) Primary and secondary metabolism, for genes that encode essential roles in plant growth, development and reproduction; (3) Structural and transport components, like transmembrane proteins, aquaporins and signal molecules; (4) Pathogenesis-related proteins (PR), for those specific to pathogen signaling and recognition and (5) Plant defense molecules, including hydrolytic proteins, responsible for degrading pathogenic structures. A table with all the statistically different expressed genes, fold-changes (FC), and functional designations by DAVID or literature is available in the supplementary material (Supplementary Table 1).

### Statistical analysis

For all analyses, the assumption of normality was checked by Shapiro–Wilk and Kolmogorov– Smirnov tests before analysis. In the case of non-normality, transformations were made based on the level of skewness, using log transformation (skewness > 1) and square root (skewness > 0.5) (Anthony et al 2021, Box and Cox 1964). Data for the two experiments were tested using Kruskal– Wallis one-way analysis of variance on ranks (p < 0.05), which was applied for significant means. Tukey’s multiple range test was applied for comparison among biomass parameters: shoot fresh Weight (SFW) and root fresh Weight (RFW). Except for the Intercellular CO_2_ (Ci), plant physiology data were subjected to square-root transformation for assimilation rate (A) and −log_2_ transformation for transpiration rate (E) and stomatal conductance (GSW) before running a mean comparison (*T*-test) to meet the assumptions of variance analysis. R-Studio 2023.12.1-402 was used for statistical analyses.

### Microbiome analysis

The platform SHAMAN was used for the metataxonomic analysis from raw reads to statistical analysis (Volant et al 2020). Sequences were clustered by the operational taxonomic unit (OTU) at 97% similarity, and OTU taxonomy was assigned using the SILVA database (McDonald et al 2012). For non-metric multidimensional scaling (NMDS), a distance matrix was generated using the Bray–Curtis distance metric in a permutational multivariate analysis of variance (PERMANOVA) test was conducted (p < 0.05) using the vegan R package (Oksanen et al 2015). The Alpha and Inverse Simpson diversity indexes were calculated using the default parameters of Volant et al. (2020). For the network analysis, the OTUs obtained from the soils as input and considering both biological associations and niche (Martins et al 2023, Poudel et al 2021), network analysis was applied to integrate microbes, plant phenotypes, and soil proprieties as a system to investigate how phytobiome structure influences crop health using the methods and codes adapted from Poudel et al (2021). Raw read data was submitted to the NCBI SRA under the accession number PRJNA1046511. Highly interactive members known as “hub” microbes, were identified. Bacterial taxa were evaluated to identify the hub microbes to determine if their association with desired plant phenotypes co-occurs more or less often than expected by chance using an OTU-OTU network. An abundance table of OTUs across different soil samples was constructed, forming the basis of the network analysis. Pairwise, Spearman’s rank correlations between the abundances of different OTUs were calculated using the rcorr function from the Hmisc package in R (Harrell Jr 2019). A significance threshold of (*p*<0.001) was applied to these correlations, which were then used to construct the adjacency matrix. Using the igraph package in R, the adjacency matrix was transformed into a network representation, with nodes representing OTUs and edges indicating significant correlations (Csardi and Nepusz 2005). Phenotype-OTU network analysis (PhONA) was constructed based on bacterial OTUs and associated disease severity as the phenotype after infection by *X. perforans* in plants. Logistic regression was utilized to establish the log odds of the probability of associations between the OTUs and the phenotype, allowing for the analysis of how each OTU’s presence or absence is related to the phenotype. The resulting significant OTUs (*p*<0.05) were transformed into a network using the igraph package in R.

## RESULTS

### The effect of antibiotic application on disease development

The application of streptomycin in the tomato rhizosphere, followed by the application of a foliar pathogen (*Xanthomonas perforans*) on the upper part of the plants, led to an increase in disease severity at 13 and 15 days after the antibiotic application of 21% and 27%, respectively (Fig1). No statistically significant difference in disease severity was found between plants that received the soil microbiome transplant (SMT) and those that did not, for any of the time points assessed (Fig. 1).

**Fig. 1.**
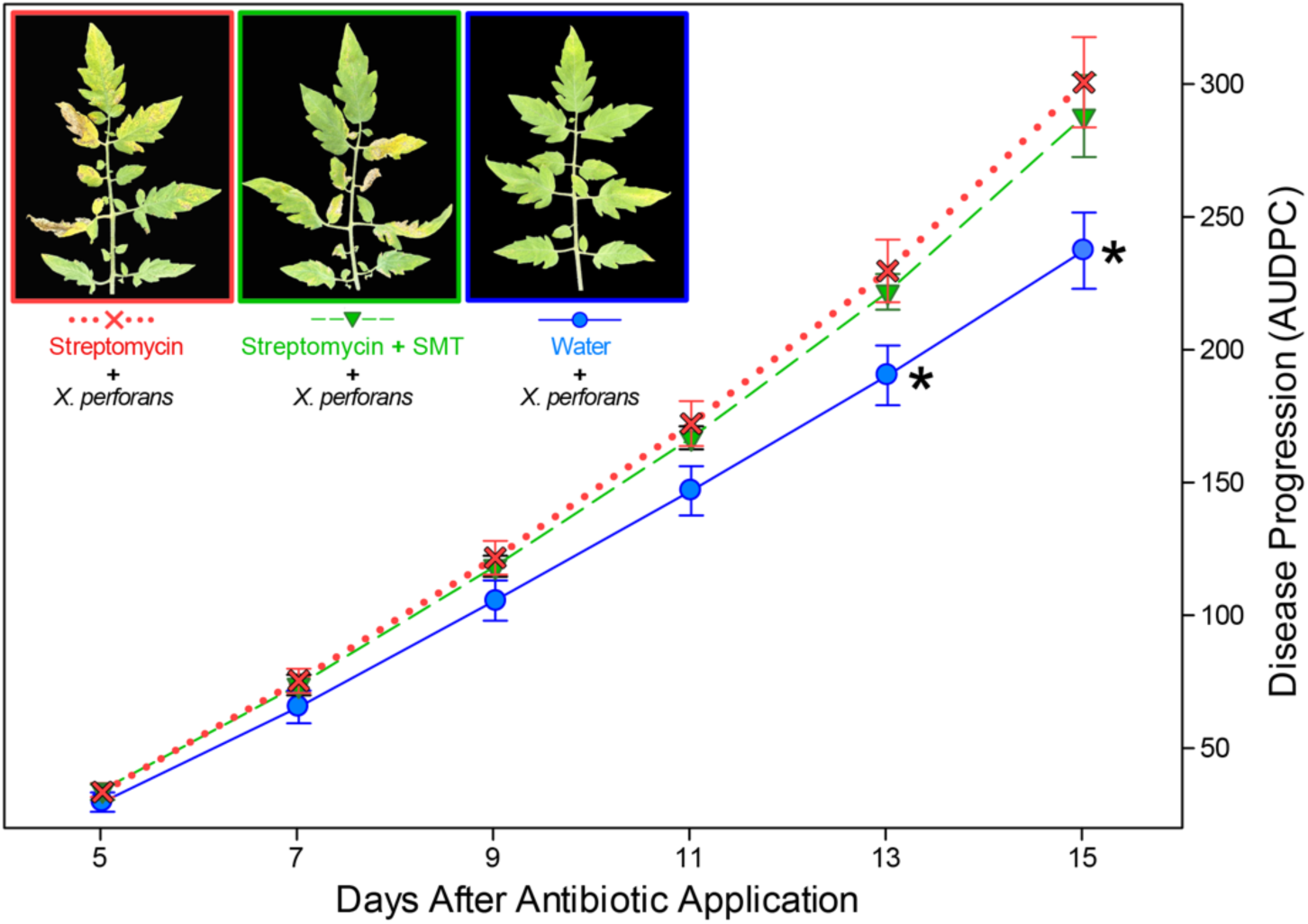
Disease progression as the area under the disease progress curve (AUDPC) of 3-week tomato seedlings cv. Alisia Craig treated with Streptomycin (250 mL, 0.6 g × L^-1^) with or without the addition of microbiome transplant (SMT) from healthy tomato plants. Control plants were treated with water. The line on each bar represents ±SE. (Means of 2 experiments and five replicates per treatment).

### Physiological & plant growth parameters

In order to investigate the physiological parameters associated with streptomycin treatment in the rhizosphere, three-time points were assessed. The first time point was a week after the antibiotic application when plants still showed no difference in disease severity among treatments. The second and third time points were at 2 and 3 weeks after antibiotic application when the plants under dysbiotic conditions started showing more disease than the negative control (Fig. 1).

The 21-day mark revealed a 0.74-fold decrease in the assimilation rate (A) for the streptomycin-only treatment, compared to control (water) (*p*<0.05; Fig. 2A).

**Fig. 2.**
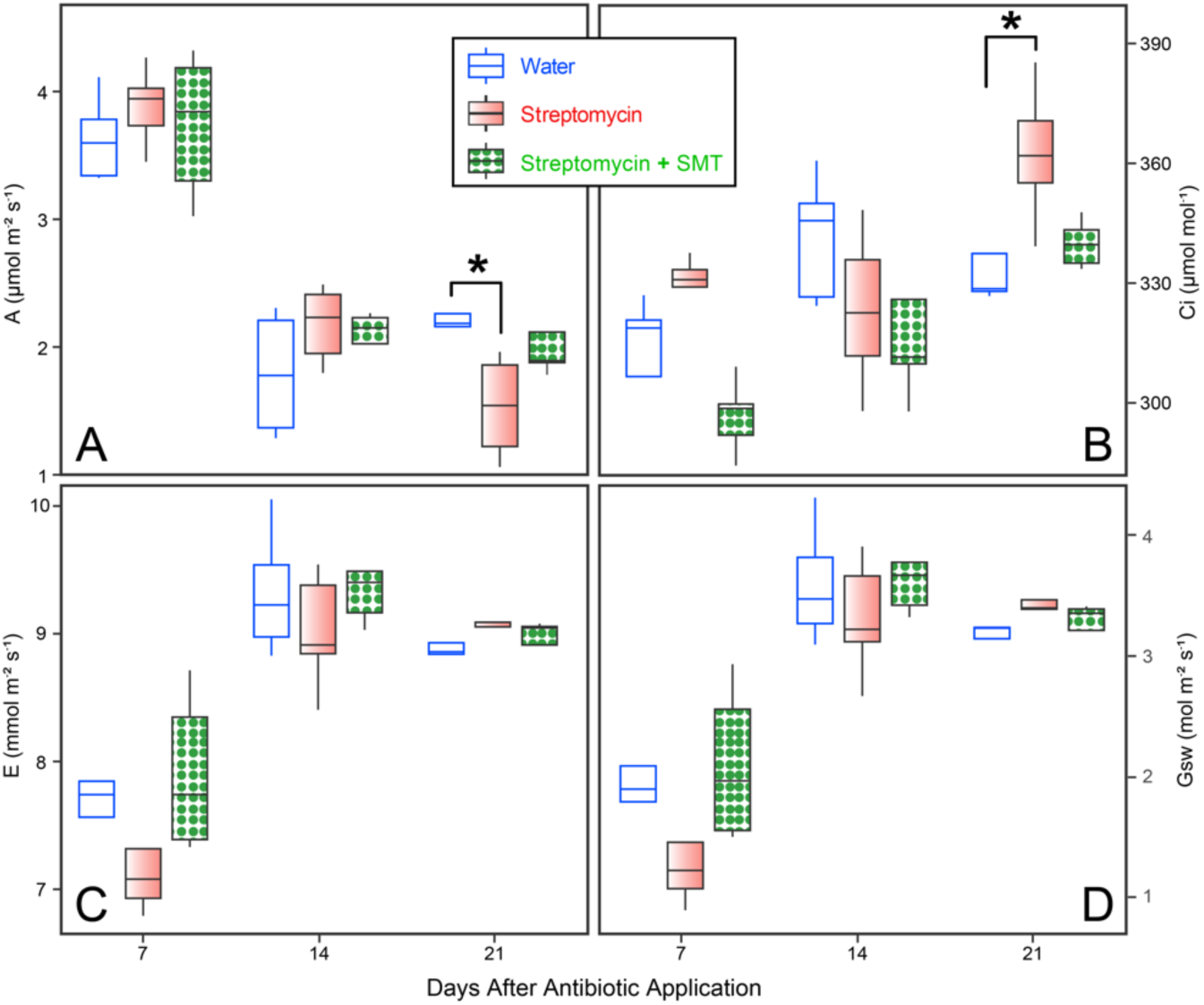
Physiological parameters evaluated in the infrared gas exchange analysis. (**A**) A = Assimilation rate (µmol m^-2^ × s^-1^); (**B**) Ci = Intercellular CO_2_ (µmol × mol^-1^); (**C**) E = Transpiration rate and (**D**) Gsw = Stomatal conductance to water vapor (mol m^-2^ × s^-1^). (Means of 2 experiments and five replicates per treatment).

Additionally, the leaf intercellular CO_2_ (Ci) levels of streptomycin-treated plants were 1.1-fold higher compared to control plants (*p*<0.05; Fig. 2B). No differences were found among the treatments for the transpiration rate (E) and stomatal conductance to water vapor (GSW) variable for the time points assessed (Fig. 2CD). Analyses of the fresh and dry weights of root and shoots (Supplementary Table 2) did not find any statistical significance among treatments.

### Rhizosphere bacterial assemblages

The top 10 most abundant phyla and genera in the rhizospheres of experimental plants can be seen in Fig. 3AB.

A significant loss of the Cyanobacteria and Patescibacteria phyla was observed when the antibiotic was applied (Fig. 3A). At a genus and phylum level, a loss in *Cylindrospermum* (Cyanobacteria)*, Conexibacter* (Actinobacteria), and *Sphingomonas* (Pseudomonata) abundance was observed. For instance, for *Cylindrospermum* a reduction of In this study, the soil microbiome transplant did not replace the loss of Cyanobacteria diversity, however, SMT enhanced the relative abundance of the Patescibacteria phylum and *Conexibacter* and *Sphingomonas* genera. The other genera and phyla did not show noticeable variation after treatment with streptomycin and attempted replacement with SMT.

**Fig. 3.**
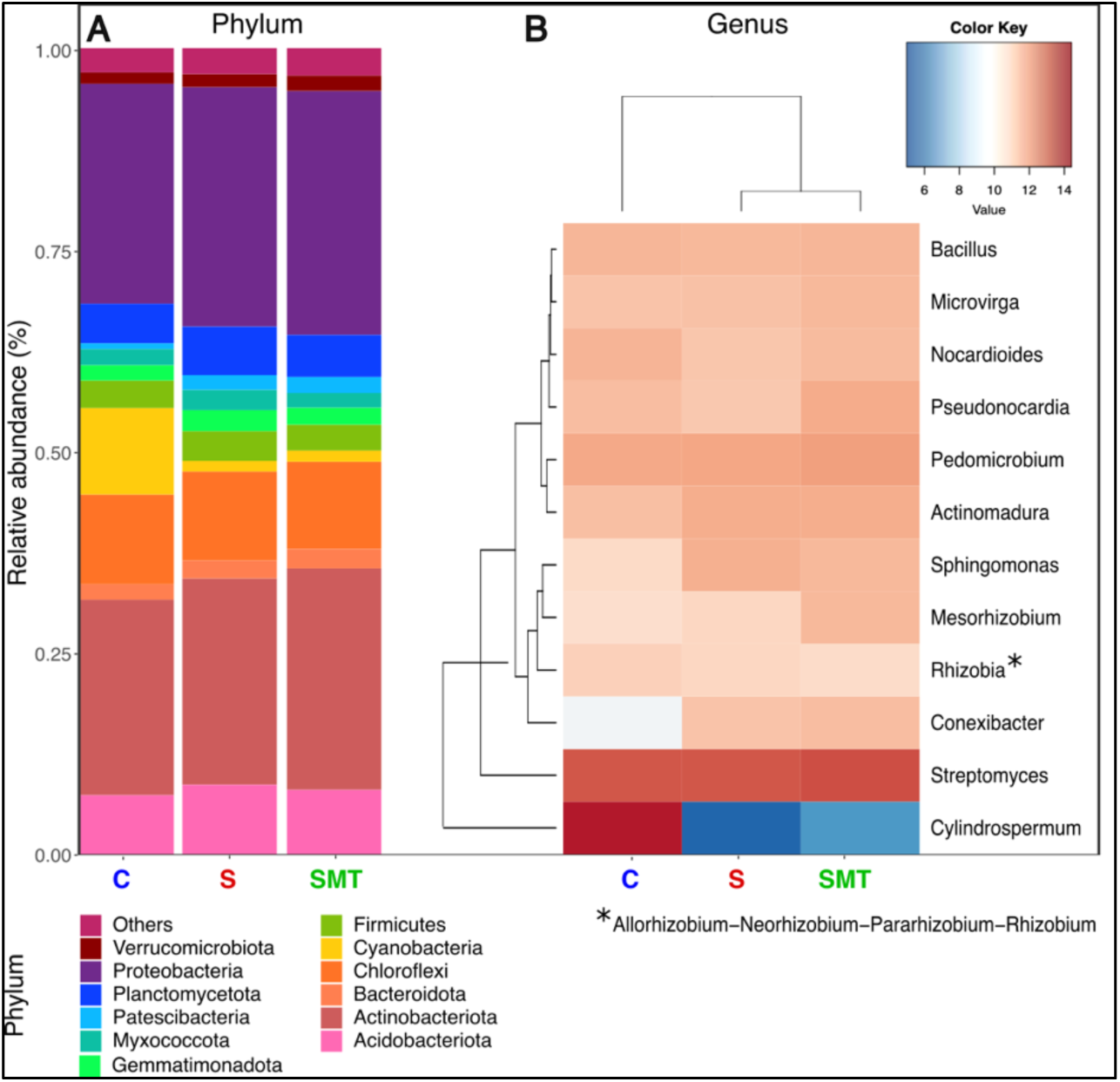
Bar plot and heatmap for the microbiome assemblages for the experiment of soil microbiome transplant (SMT) after treatment with Streptomycin (S) and water used as control (C). (**A**) Bar plot at phylum level with the ten most abundant with labels organized according to the relative abundance. (**B**) Heatmap of relative abundance at a genus level representing the most abundant features of each treatment. (Means of 2 experiments and five replicates per treatment).

Streptomycin treatment presented a higher alpha microbial diversity at the phylum level compared to SMT treatment (*p*-value = 0.036; Fig. 4A), indicating a slightly higher average of phylum among samples of the same treatment. Additionally, soil microbiome transfer (SMT) treatment at a genus level uses the Inverse Simpson index to assess evenness, also showing increase (*p*-value = 0.035), indicating reduction in the counts of previously dominant genera when compared to the water-only treatment. The non-metric multidimensional scaling (NMDS) plot revealed distinct patterns among three treatments (*p*=0.02; Fig. 4C), also showing difference in clusters of the two replicas of the experiment.

**Fig. 4.**
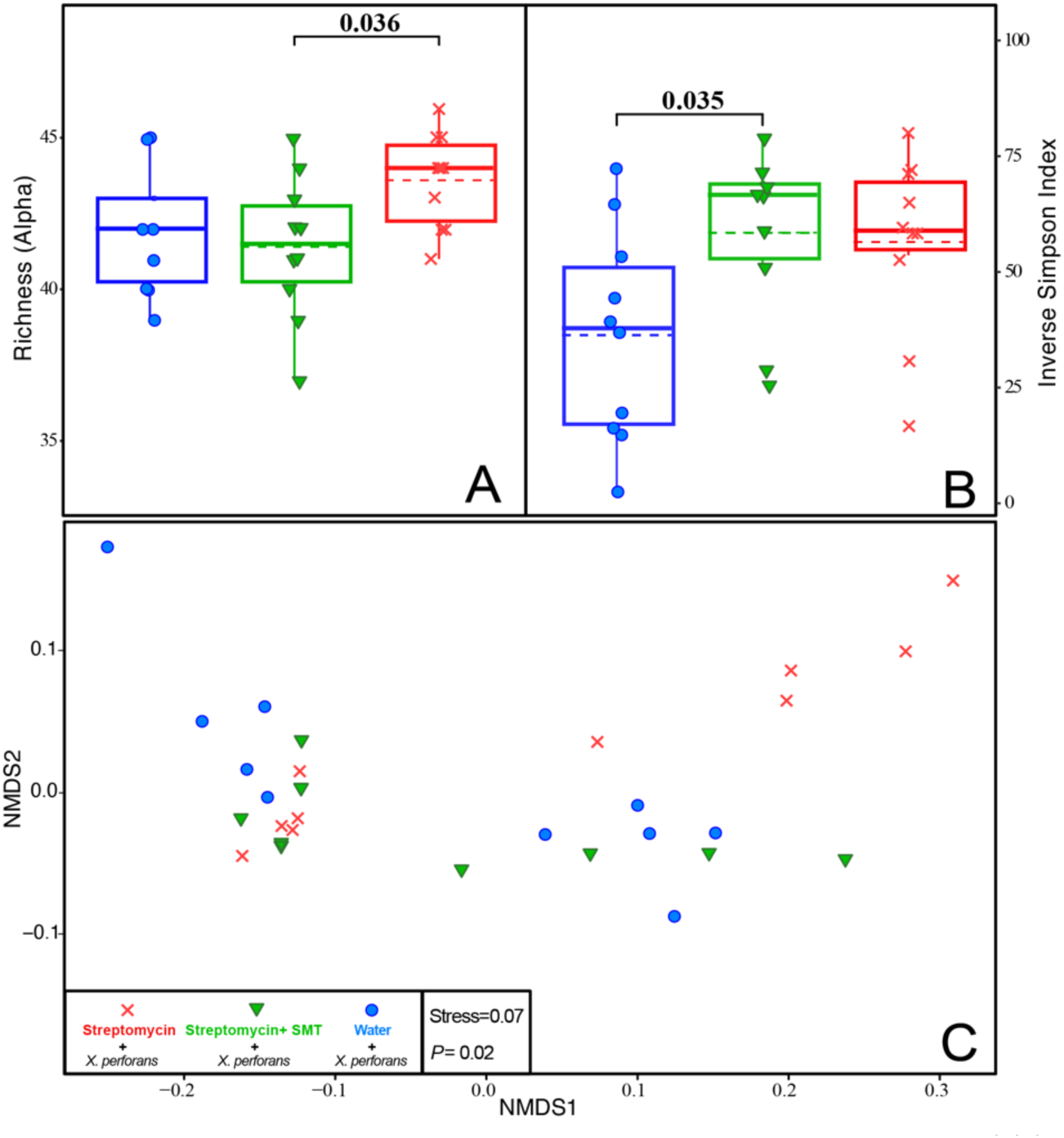
Ordination and **e**cological indexes for the experiment of soil microbiome transplant (SMT) after treatment with Streptomycin (S), where water was used as control (C). (**A**) Alpha Richness at a phylum level. (**B**) Inverse Simpson diversity at a genus level. (**C**) A non-metric multidimensional scaling (NMDS) of global 16S rRNA gene composition for the control (water) treatments with Streptomycin with (SMT) and without (S) soil microbiome transplant. (Means of 2 experiments and five replicates per treatment).

Moreover, in both replicas the Water group (control) clustered distant from the streptomycin-treated group, suggesting a distinct rhizosphere microbiome composition or proportion. On the other hand, the SMT treatment overlapped with the negative control. The low-stress value suggests a reliable representation of these differences in the microbial communities.

### Operational taxonomic units (OTU)-phenotype (disease resistance) networks

In the OTU-OTU networks (Fig. 5-ABC), the results illustrated significantly higher relative abundances of *Cylindrospermum* in the water-only group (Fig. 5-A) compared to the soil microbiome transplant and streptomycin treatment groups, where *Streptomyces* emerged as the predominant organism.

**Fig. 5.**
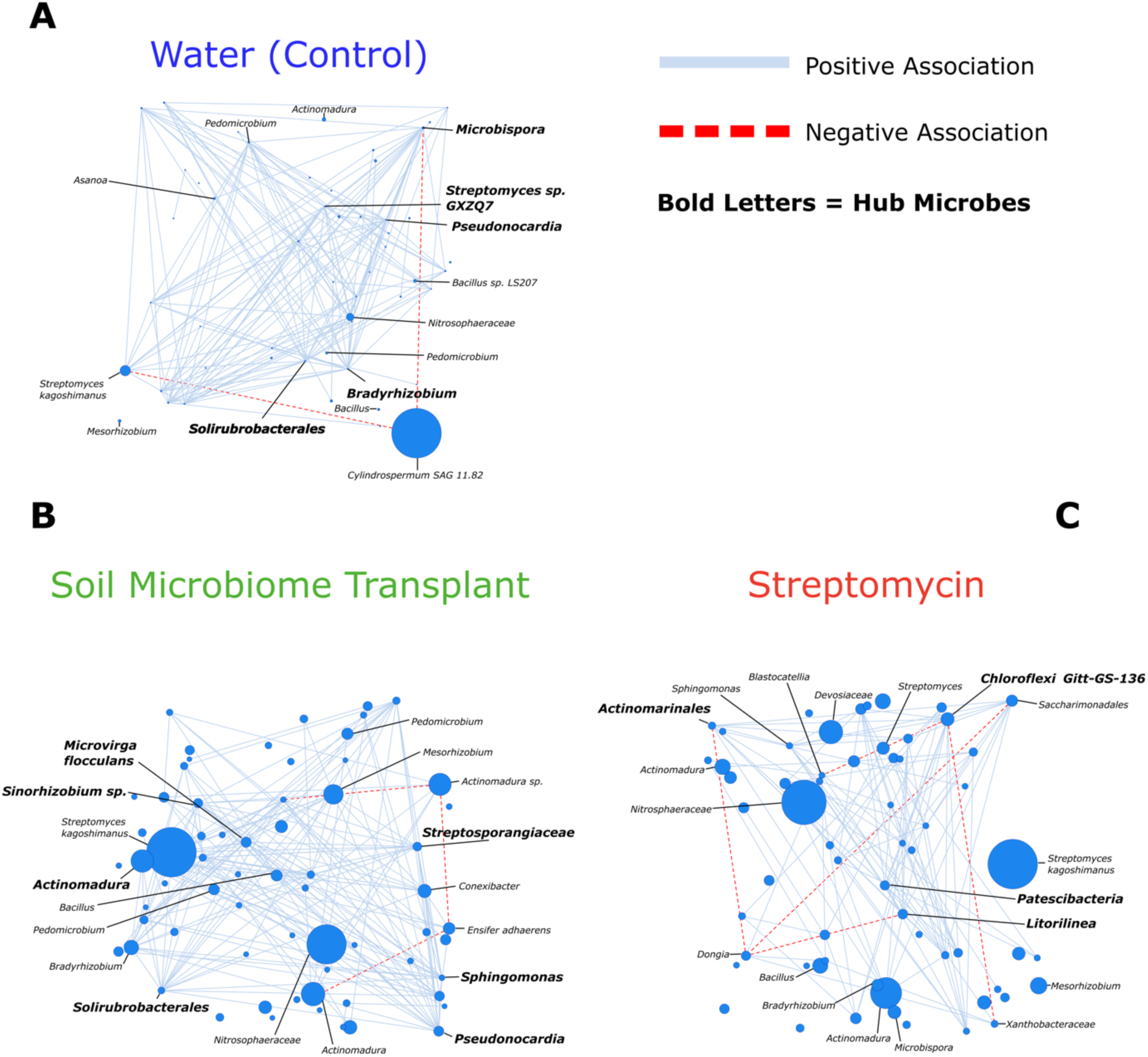
OTU-OTU networks pictured represent each treatment type, with node size corresponding to relative abundance and edge color corresponding to positive (green) and negative (red) correlations within the three treatments: water-only, used as negative control (C), streptomycin-treated soil (S) and soil microbiome transplant in a previously streptomycin-treated soil (SMT) (Means of 2 experiments and five replicates per treatment). Hub microbes, defined as the nodes in the top 10% of degree values, are highlighted with a bold font.

The OTU-OTU network analysis revealed the soil microbiome transplant treatment showed the overall highest connectivity (174), followed by water-only (151) and streptomycin (105) (Table 1).

**Table 1.** Summary of Key Network Metrics for Microbial Community Analysis.

| Metric | Water | Soil Microbiome Transfer | Streptomycin |
| --- | --- | --- | --- |
| Number of Nodes<sup>a</sup> | 52 | 65 | 61 |
| Total Number of Connections<sup>b</sup> | 151 | 174 | 105 |
| Positive edges | 149 | 171 | 100 |
| Negative edges | 2 | 3 | 5 |
| Average Degree | 5.81 | 5.35 | 3.44 |

<sup>a</sup>Number of nodes established with an abundance threshold of >5000 OTU counts.
<sup>b</sup>Total number of connections (edges) established with a p-value < 0.001.

Additionally, the water-only group (C) showed the highest overall average degree compared to the soil microbiome transfer (SMT) and streptomycin (S) treatments.

In the OTU-Phenotype network analysis, the top 35 genera of bacteria were identified based on statistical significance (p < 0.05) in association with the phenotype ‘disease severity,’ as derived from logistic regression analysis. Among these genera, the majority belonged to the phylum Proteobacteria, with 23 nodes, followed by Actinobacteria with 5 nodes (Fig. 6). Moreover, organisms such as *Bauldia*, *Roseiflexaceae*, and *Hirschia* are positively associated with healthier plants (lower disease severity), while *Nordella*, *Devosiaceae*, and *Pedomicrobium* are associated with disease progress (higher disease severity) (p<0.05; Fig. 6).

**Fig. 6.**
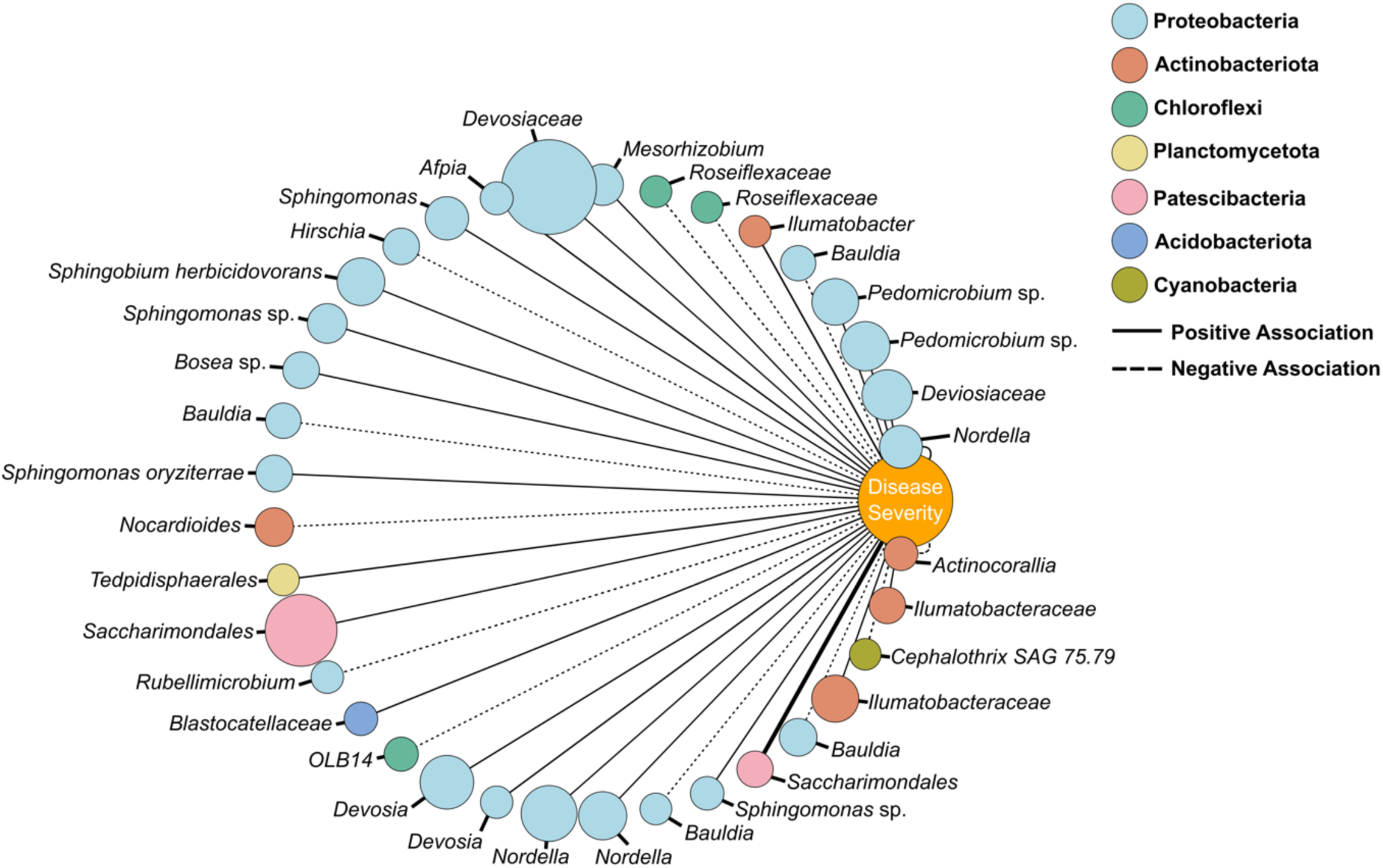
The OTU-phenotype network displays 35 significant (p<0.05) OTUs (Operational Taxonomic Units) associated with the log odds of the probability (thorough a logistic regression analysis) of either higher or lower disease severity. The node color corresponds to the respective phylum of the OTU, and node size is according to relative abundance. Edge width is based on the strength of the probability coefficient, with positive and negative associations displayed by solid and dotted lines, respectively. (Means of 2 experiments and five replicates per treatment).

### Transcriptome results

Of all the genes evaluated, 113 were statistically different in the streptomycin treated plants (P≤0.05) in terms of expression compared to the Water-only treatment, with 76 up-regulated and 37 down-regulated. Among the statistically different genes, 65 could be assigned a functional group using the DAVID clustering tool, while 48 were manually grouped (Fig. 7-A).

**Fig. 7.**
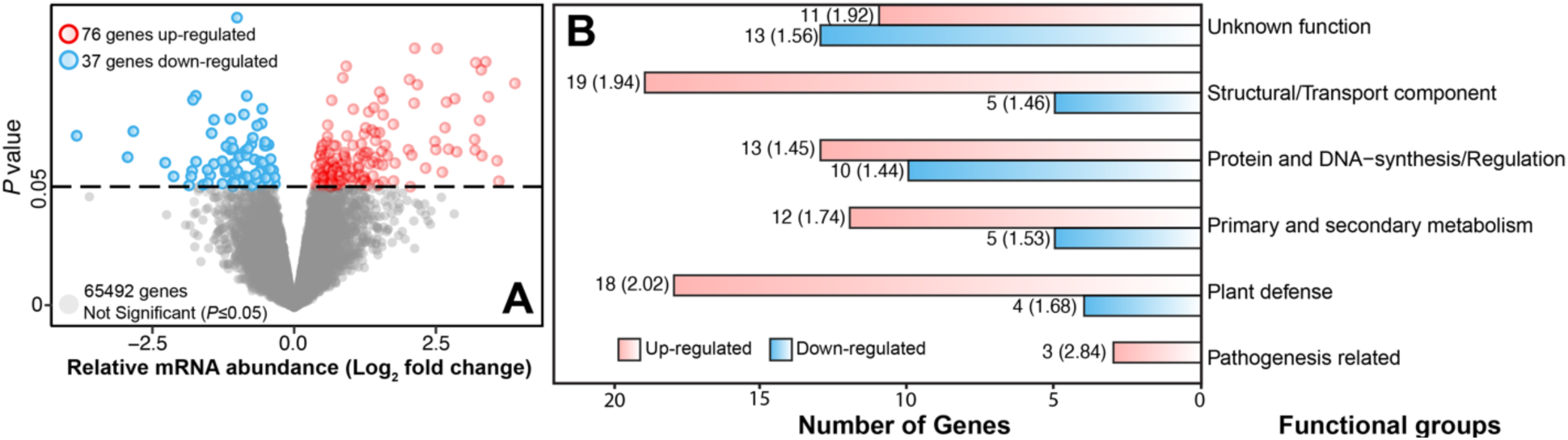
Transcriptome results, and functional grouping of differentially expressed genes of 3-week-old tomato plants cv. Alisia Craig was assessed 24 h after pathogen (*Xanthomonas perforans*) inoculation. (**A**) Volcano plot with the distribution of the relative abundance of quantified mRNA and expresses them in Log2 of fold change, which describes how much a measurement changes between an original and a secondary measurement, in this case, control and treatment with. Statistically different (*p<*0.05) according to False Discovery Rate (FDR) test; up-regulated (blue) and downregulated (red) genes are expressed based on the fold-change with respective counts; (**B**) Clustering of genes based on biological function. Up-regulated (blue) and downregulated (red) bars show the number of genes assigned to each function and the mean value for the fold change in streptomycin treated plants compared to water treated plants. (Means of 2 experiments and five replicates per treatment).

To provide a dimension of the possible biological functions that may be modified due to changes in the dynamics of the root microbiome, the genes were grouped based on their biological functions (Fig. 7-B) when known. For structural components and those responsible for transporting molecules, 19 genes were up-regulated, and five were downregulated, with averages for Log_2_ fold change of 1.94 and 1.46, respectively.

Genes related to synthesizing and regulating genetic material, proteins, and up-and down-regulated genes were paired with 13 and 10 genes and 1.45 and 1.44 of Log_2_ FC, respectively. Genes directly or indirectly linked to plant defense showed a more significant discrepancy, with hyperregulation of 18 genes and an average Log_2_ FC of 2.02. In comparison, the reduction in expression was only observed in 4 genes, with an average Log_2_ FC of 1.68. Proteins related to pathogenesis, essential in activating the plant’s immune system and its interactions with pathogenic organisms, were only up-regulated, with three genes detected and an average Log_2_ FC of 2.84.

## DISCUSSION

This study found that inducing an imbalance in the rhizosphere microbiome, also known as dysbiosis, has significant effects on the infection and progression of above-ground plant diseases, as is shown for bacterial spot disease caused by *Xanthomonas perforans*, with lasting effects even after trying to replace the microbiome disrupted by the application of streptomycin. Although no health recovery was observed after the soil microbiome transplant (SMT) in plants subjected to streptomycin while infected with *X. perforans*, a slight difference in disease progress over time was observed. At 15 days after antibiotic application, a difference of 4% in disease severity was found between streptomycin-treated and the SMT treatment (Fig. 1).

The impact of soil and rhizosphere microbiome on soilborne diseases in plants is well-established (Lee et al 2021, Pasche et al 2023), demonstrating the significant role of below-ground microbial communities in plant health and disease resistance. However, our understanding of how these soil-root microbiome dynamics influence above-ground diseases, particularly foliar diseases, is still underexplored (Trivedi et al 2020). In this study, the disruption in the root microbiome induced by the application of streptomycin led to an increase in tomato bacterial spot disease caused by *Xanthomonas perforans*. This trend was also observed in studies where the disruption in the rhizosphere microbiome caused by Vancomycin (500 µg × mL^-1^) also led to higher severity in bacterial wilt disease caused by *Ralstonia solanacearum* (Lee et al 2021). Although no significant reduction in the disease severity was observed for the SMT treatment when applied to plants under dysbiosis (subjected to streptomycin), it is possible that the timing of application was not early enough (only 24 h after the pathogen application) to trigger plant immunity and offer plant protection as shown in other studies (Armstrong et al 2022, Dexter et al 2018). Since foliar colonization can induce microbial recruitment of beneficial microorganisms in the roots (Dastogeer et al 2022), the presence of antibiotic residues on the soil can impact plant’s recruitment of microbes capable of aiding in above-ground plant health (Berendsen et al 2018, Cycoń et al 2019, Trivedi et al 2020, Trivedi et al 2022).

Plant health is complex, with many factors contributing to the final disease severity, including physiological indexes (Trivedi et al 2020). The physiological parameters measured pointed to a lower CO_2_ assimilation in the tomato plants under dysbiosis, accompanied by higher intercellular CO_2_ concentrations, suggesting that the plant is not efficiently converting CO_2_ into sugars through photosynthesis, indicating a possible stress (Ethier et al 2006). Photosynthesis can be intimately influenced by beneficial plant-microbe interactions, primarily through plant hormone production and aids in nutrient acquisition (Lee et al 2022, Martins et al 2015).

After natural root microbiome dysbiosis inducted by streptomycin application, a drastic reduction in Cyanobacteria was observed, a phylum representing a promising agricultural input due to its plant growth-promoting capabilities, including but not exclusive to tomato plants (Lee et al 2022, Rana et al 2012, Toribio et al 2022). The disruption in Cyanobacteria in the microbiome assemblages was also extended to the genus level, with a decrease in *Cylindrospermum* relative abundance across all streptomicyn-treated samples, a known plant growth-promoting Cyanobacteria capable of induced physiological changes, as observed in lupin bean (Haroun and Hussein 2003, Kollmen and Strieth 2022). OTU network analysis revealed that *Cylindrospermum* showed a higher relative abundance in the negative control compared to the treatments that received antibiotics, highlighting its sensitivity and negative association with *Streptomyces*, the dominant organism in both antibiotic-treated groups (Santini et al 2021). The transplant of soil microbiome after exposure to streptomycin was not able to restore noticeable levels of Cyanobacteria, possibly due to the significant sensitivity of Cyanobacteria, including *Cylindrospermum,* to streptomycin even in concentrations as low as 0.5 µg × mL^-1^ (Mishra et al 2013). Re-composition of previous microbiome structures through soil microbiome transplant can be difficult. Even though plant growth promotion can be consistently noticed as a lasting beneficial effect due to the increased support of microbial activity and ecological functions, the actual replacement of the disrupted microbial community only takes place in 16% of the trials (Yergeau et al 2015). *Conexibacter* and *Sphingomonas* significantly increased relative abundance at the genus level after soil exposure to streptomycin. *Conexibacter* and *Sphingomonas* have been reported as antibiotic-resistant, including to streptomycin (Vanbroekhoven et al 2004, Zhao et al 2019). Streptomycin can remain undegraded in agricultural soils for two days, with the dissipation of 50% of the parent compound in about six days at room temperature (Demars et al 2024, Klein and Pramer 1961). Unexpectedly, in this study, plants that received streptomycin presented an increase in alpha diversity, which is the average number of phyla among samples of the same treatment (Pielou 1966, Walters and Martiny 2020). The higher alpha diversity could result from the streptomycin reducing or eliminating dominant phyla and, therefore, increasing the abundance of bacteria taxa in lower populations (Freestone and Inouye 2006, Walters and Martiny 2020). Similarly, the increase in the Simpson index at a genus level for the treatments that had contact with streptomycin (S and SMT) (Fig. 4B) may signal a reduction of highly dominant genera. Antibiotics might suppress dominant (high relative abundance) but sensitive microbial taxa (i.e., *Cylindrospermum*) (Fig. 3), allowing less abundant resistant species to flourish and cause a reduction in the Simpson index, which captures the evenness of relative abundances among the taxa and can signal the presence of dominant species in the samples (Hillebrand et al 2018, Le Page et al 2019, Simpson 1949).

The transcriptome results suggest that subjecting rhizosphere-associated soil to streptomycin doses can cause significant changes in plant gene expression. The majority of genes significantly altered still have no specific function designation, followed by genes related to the physical structure of the cell and transport mechanisms, which were both up- and down-regulated. These results suggest that the plants, after infection, were remodeling their cellular components and alterations in nutrient or molecular transport in response to the treatment (Bai et al 2022, Mukai et al 2022). Pathogen-induced gene changes associated with core biological processes, such as protein synthesis and gene regulation, could impact the plant’s overall metabolism and growth due to an expressive physiological cost in detecting and suppressing pathogens (Heil 2001, Yu et al 2022). The upregulation of genes related to primary and secondary metabolism may reflect an increase in metabolic activities due to the activation of routes of systemic acquired resistance as defense mechanisms or a more costly growth to compensate for pathogen stress and damage (Qu et al 2020, Yu et al 2022). An apparent increase in genes related to plant defense points to the activation of the plant’s immune response, possibly to counteract the adverse effects of dysbiosis and pathogen attack (Arnault et al 2023, Poppeliers et al 2023, Teixeira et al 2021). An up-regulation of the pathogenesis-related genes indicates that the plant may be increasing its defense mechanisms against potential pathogens, which could be more active or prevalent due to the changes in the soil microbiome caused by the antibiotic application (Agri et al 2022, Chinnappan et al 2018).

This study sheds light on the impacts of rhizosphere dysbiosis on above-ground plant health, using tomato plants and *Xanthomonas perforans* as a pathosystem. It also presented molecular, physiological, and microbiological explanations for reduced plant defense under dysbiosis. It was observed that antibiotic treatment aimed at disrupting the soil microbiome led to an increase in bacterial spot disease severity. However, attempts to restore the microbiome composition and balance through soil microbiome transplants (SMT) yielded some mitigation of disease progression, underscoring the potential of microbiome management in plant disease control, but no evident success in restoring the original microbiome diversity. The reduction in beneficial microbial taxa, particularly Cyanobacteria (*Cylindrospermum*), alongside alterations in plant gene expression, highlights the intricate interplay between soil microbial communities and plant health. The findings advocate for a deeper understanding of soil microbiome dynamics and their implications for sustainable agricultural practices, emphasizing the need for strategies that preserve or restore beneficial microbial communities to bolster plant resilience against pathogens.

## Acknowledgments

This work was supported by the USDA National Institute of Food and Agriculture (NIFA) project no. 2022-68015-36721 and Hatch project no. 1024881. The authors thank Dr. Jeff Jones, who kindly shared the bacterial strain *Xanthomonas perforans* XP1-6 used in this study.

