## Supplementary Table 1 and Table 2 for "The Underground World of Plant Disease: How Does Rhizosphere Dysbiosis Affect Plant Health Above-ground?"

| **Supplementary Table 2**. Morphometric analysis of plants from the three treatments. | | | | | | | | | |
| --- | --- | --- | --- | --- | --- | --- | --- | --- | --- |
|  |  | Roots | | | | Shoots | | | |
|  | Treatment | *Fresh Weight (g) | | Dry Weight (g) | | Fresh Weight (g) | | Dry Weight (g) | |
|  | C | 3.179 | a | 0.463 | a | 19.116 | a | 2.846 | a |
|  | S | 3.095 | a | 0.483 | a | 24.196 | a | 3.602 | a |
|  | SMT | 3.487 | a | 0.490 | a | 20.016 | a | 2.984 | a |
| *Means followed by the same letter in the column do not differ from each other in a Tukey test at 5% probability. | | | | | | | | | |

| **Supplementary Table 1.** Chemical parameters assessment for the soil used in this study. Chemical parameters used in soil fertility evaluation. Macronutrients, micronutrients, pH, and organic matter (OM). | | | | | | | | | | |
| --- | --- | --- | --- | --- | --- | --- | --- | --- | --- | --- |
| NO_3_^-^ | Cu^2+^ | Mn^2+^ | Zn^2+^ | NH_4_^+^ | Ca^2+^ | Mg^2+^ | K^+^ | P^3-^ | pH | OM (%) |
| mg × kg^-1^ in the soil | | | | | | | | | 6.85 | 6.79 |
| 3.71 | 2.94 | 32.51 | 24.65 | 3.99 | 2722.07 | 422.54 | 180.49 | 569.76 |  |  |
